# Oxidative pathways of deoxyribose and deoxyribonate catabolism

**DOI:** 10.1101/205583

**Authors:** Morgan N. Price, Jayashree Ray, Anthony T. Iavarone, Hans K. Carlson, Elizabeth M. Ryan, Rex R. Malmstrom, Adam P. Arkin, Adam M. Deutschbauer

**Affiliations:** Environmental Genomics and Systems Biology, Lawrence Berkeley National Laboratory, Berkeley CA, USA; QB3/Chemistry Mass Spectrometry Facility, University of California, Berkeley CA, USA; DOE Joint Genome Institute, Walnut Creek CA, USA; Department of Bioengineering, University of California, Berkeley CA, USA; Department of Plant and Microbial Biology, University of California, Berkeley CA, USA

## Abstract

Using genome-wide mutant fitness assays in diverse bacteria, we identified novel oxidative pathways for the catabolism of 2-deoxy-D-ribose and 2-deoxy-D-ribonate. We propose that deoxyribose is oxidized to deoxyribonate, oxidized to ketodeoxyribonate, and cleaved to acetyl-CoA and glyceryl-CoA. We have genetic evidence for this pathway in three genera of bacteria, and we confirmed the oxidation of deoxyribose to ketodeoxyribonate *in vitro*. In *Pseudomonas simiae*, the expression of enzymes in the pathway is induced by deoxyribose or deoxyribonate, while in *Paraburkholderia bryophila* and in *Burkholderia phytofirmans*, the pathway proceeds in parallel with the known deoxyribose 5-phosphate aldolase pathway. We identified another oxidative pathway for the catabolism of deoxyribonate, with acyl-CoA intermediates, in *Klebsiella michiganensis*. Of these four bacteria, only *P. simiae* relies entirely on an oxidative pathway to consume deoxyribose. The deoxyribose dehydrogenase of *P. simiae* is either non-specific or evolved recently, as this enzyme is very similar to a novel vanillin dehydrogenase from *Pseudomonas putida* that we identified. So, we propose that these oxidative pathways evolved primarily to consume deoxyribonate, which is a waste product of metabolism.

**Importance:** Deoxyribose is one of the building blocks of DNA and is released when cells die and their DNA degrades. We identified a bacterium that can grow with deoxyribose as its sole source of carbon even though its genome does not encode any of the known genes for breaking down deoxyribose. By growing many mutants of this bacterium together on deoxyribose and using DNA sequencing to measure the change in the mutants’ abundance, we identified multiple protein-coding genes that are required for growth on deoxyribose. Based on the similarity of these proteins to enzymes of known function, we propose a 6-step pathway in which deoxyribose is oxidized and then cleaved. Diverse bacteria use a portion of this pathway to break down a related compound, deoxyribonate, which is a waste product of human metabolism and is present in urine. Our study illustrates the utility of large-scale bacterial genetics to identify previously unknown metabolic pathways.

## Introduction

Deoxyribonucleic acid (DNA) is present in every cell, and once released into the environment by cell lysis, some of the DNA is probably hydrolyzed to release 2-deoxy-D-ribose (Levy-Booth et al. 2007). Indeed, some nucleosidases are specific for 2’-deoxyribonucleosides over ribonucleosides and release deoxyribose (Koszalka and Krenitsky 1979). The only previously-known pathway for catabolizing deoxyribose (of which we are aware) involves deoxyribose kinase (*deoK)* (Tourneux et al. 2000) and deoxyribose 5-phosphate aldolase (*deoC)* (Valentin-Hansen et al. 1982), which yields acetaldehyde and D-glyceraldehyde 3- phosphate (Figure 1A).

During a screen for the growth of *Pseudomonas simiae* WCS417 on various carbon sources (Price et al. 2018), we found that it grows on deoxyribose even though its genome does not contain either *deoK* or *deoC*. Using a pool of over 100,000 randomly-barcoded transposon mutants of *P. simiae*, we tested which genes are important for growth in 89 different conditions, including growth with deoxyribose as the sole source of carbon (Price et al. 2018).

Here, we use this genetic data to identify a novel oxidative pathway for deoxyribose catabolism. An intermediate in the proposed pathway is 2-deoxy-D-ribonate, which is a waste product of various metabolisms (Kappen and Goldberg 1989;Lerma-Ortiz et al. 2016) and has been detected in human urine (Chamberlin and Sweeley 1987). We found that *P. simiae* uses many of the same genes to catabolize deoxyribonate as deoxyribose, and that these genes are induced during growth on either deoxyribose or deoxyribonate. We also reconstituted the enzymatic conversion of deoxyribose to deoxyribonate to ketodeoxyribonate *in vitro.*

We then used pooled mutant fitness assays to study the catabolism of deoxyribose and deoxyribonate in other bacteria. In *Paraburkholderia bryophila* 376MFSha3.1 and in *Burkholderia phytofirmans* PsJN, similar genes as in *P. simiae* are involved in the catabolism of both deoxyribose and deoxyribonate. *Klebsiella michiganensis* M5al uses an analogous pathway with acyl-CoA intermediates to consume deoxyribonate (but not deoxyribose). Thus, diverse bacteria use oxidative pathways to consume deoxyribose and deoxyribonate.

## Results

### The genetic basis of growth on deoxyribose and deoxyribonate in *Pseudomonas simiae*

To supplement the previously-described mutant fitness data for *P. simiae* WCS417 (Price et al. 2018), we tested deoxyribose as the carbon source on two additional days, and we also tested deoxyribonate as the carbon source. For each experiment, we grew a library of over 100,000 barcoded mutant strains together and used DNA sequencing to measure the change in each strain’s abundance. Overall, our dataset for *P. simiae* included 6 genome-wide assays of fitness in deoxyribose, 3 assays of fitness in deoxyribonate, and 153 other fitness assays from diverse conditions, including 49 other carbon sources.

From these data, we identified 13 genes that were consistently important for utilizing deoxyribose and were not important for fitness in most other conditions (see Methods). The fitness data for these genes is shown in Figure 1B: each fitness value is the log_2_ change of the relative abundance of mutants in that gene during the experiment. In the genome of *P. simiae*, 11 of the 13 genes cluster into three groups of nearby genes that are on the same strand and are probably operons.

Of the 13 genes, eight are likely catabolic enzymes:

- two subunits of a molybdenum-dependent dehydrogenase (PS417_10890, PS417_10885) that are distantly related to isoquinoline 1-oxidoreductase *iorAB* ((Lehmann et al. 1995); 49% or 29% amino acid identity)
- a cytochrome c that is in the same putative operon (PS417_10880) and might accept electrons from the *iorAB*-like dehydrogenase
- a dehydrogenase of the short chain dehydrogenase/reductase (SDR) superfamily (PS417_07245)
- a β-keto acid cleavage enzyme (PS417_07250) (Bastard et al. 2014)
- a lactonase (PS417_07255)
- glycerate kinase (PS417_13970)
- and acetyl-CoA C-acetyltransferase (PS417_10515).

The remaining genes that are specifically important for deoxyribose utilization are two putative transporters, two regulatory genes, and *mobA* (PS417_21490), which is involved in the biosynthesis of molybdenum cofactor and is probably required for the activity of the molybdenum-dependent dehydrogenase (PS417_10890:PS417_10885). Most of these genes are also important for growth on deoxyribonate (Figure 1B). The exceptions are the two subunits of the molybdenum-dependent dehydrogenase, the associated cytochrome c, *mobA*, and perhaps the lactonase (Figure 1B); we therefore propose that these genes act upstream of deoxyribonate formation (Figure 1C).

**Figure 1:**
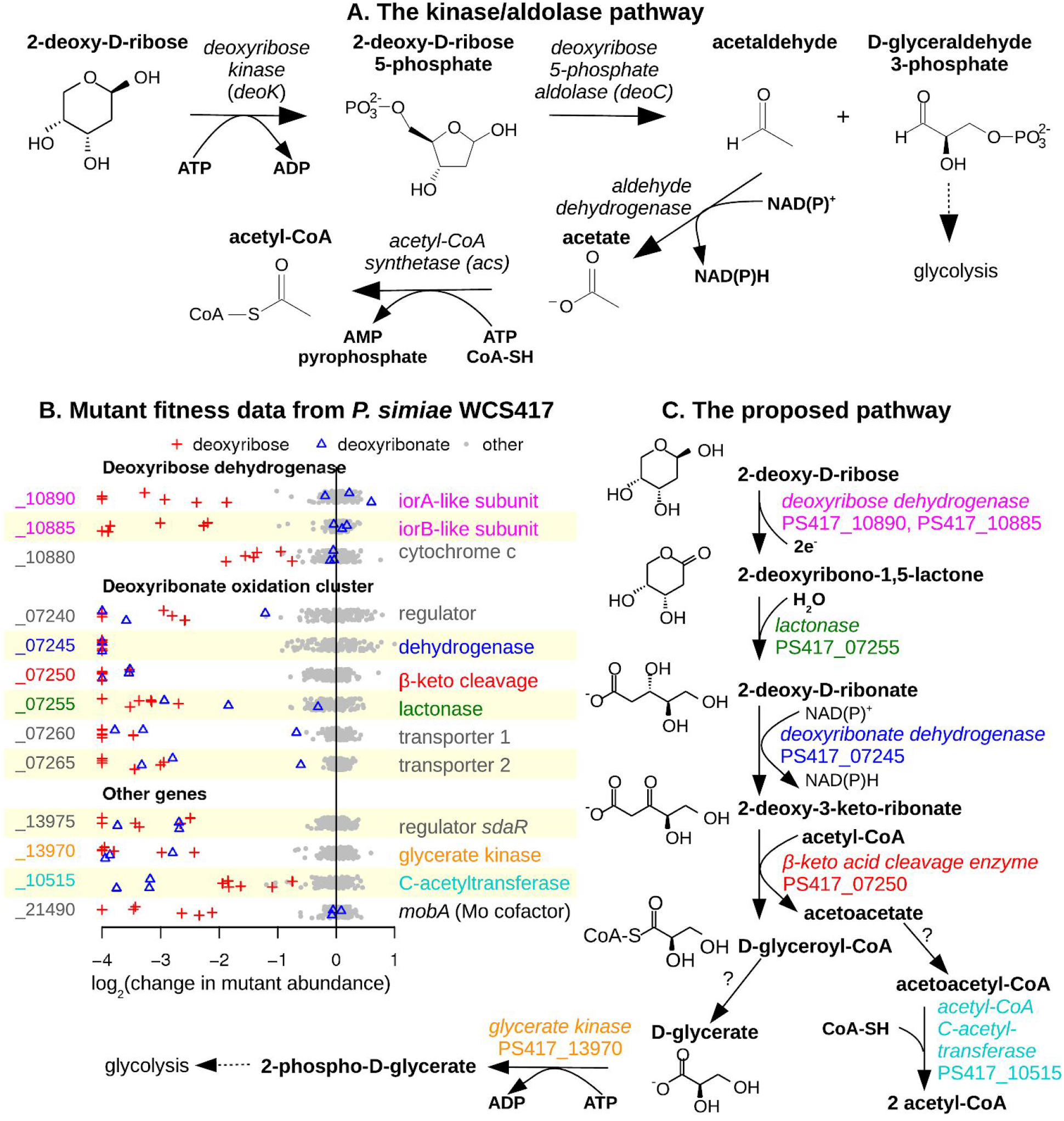
An oxidative pathway for deoxyribose utilization in *Pseudomonas simiae* WCS417. (A) The previously-described kinase/aldolase pathway. (B) Gene fitness in *P. simiae*. Each point shows the gene’s fitness value (*x* axis) in a different experiment. Fitness values under −1 indicate a defect in growth. Values outside of the plot’s range are shown at −4 or +1. Within each gene’s section, the y axis is random. Experiments with deoxyribose or deoxyribonate as the carbon source are indicated by symbols. (C) The proposed pathway for deoxyribose catabolism in *P. simiae*. Putative enzymes are color coded to match the gene descriptions in panel B.

To confirm the genome-wide fitness data, we obtained individual mutant strains for some of the key genes and grew them with either D-glucose, deoxyribose, or deoxyribonate as the sole source of carbon. We found that a transposon mutant of the *iorB*-like subunit of the molybdenum-dependent dehydrogenase (PS417_10885) grows on glucose or deoxyribonate but not on deoxyribose (Figure S1). Mutants of the SDR dehydrogenase (PS417_07245) or the β-keto acid cleavage enzyme (PS417_07250) grow on glucose but not on deoxyribose or deoxyribonate (Figure S1).

### The proposed pathway for deoxyribose catabolism in *P. simiae*

The genetic data suggest that *P. simiae* WCS417 catabolizes deoxyribose by an oxidative pathway (Figure 1C). Because the molybdenum-dependent dehydrogenase was not required for growth on deoxyribonate, and because the lactonase was more important for growth on deoxyribose than on deoxyribonate (Figure 1B), it appears that the molybdenum-dependent dehydrogenase oxidizes deoxyribose to a lactone, which is hydrolyzed to deoxyribonate by the lactonase. We tentatively predict that the lactone intermediate is 2-deoxy-D-ribono-1,5-lactone instead of a 1,4-lactone because the pyranose form of deoxyribose is more stable in solution at 30°C (Lemieux et al. 1971).

Oxidation of the deoxyribonate by the SDR enzyme could yield 2-deoxy-3-keto-D-ribonate, which is a β-keto acid and could be cleaved by the β-keto acid cleavage enzyme (PS417_07250). β-keto acid cleavage enzymes remove an acetyl group from a β-keto acid and transfer the acetyl group to acetyl-CoA (Bastard et al. 2014). In the proposed pathway, the β-keto acid cleavage enzyme is expected to yield D-glyceryl-CoA and acetoacetate. The requirement for the glycerate kinase suggests that the D-glyceryl-CoA is converted to glycerate. We looked for a thiolase or CoA-transferase that might act on D-glyceryl-CoA and that is important for deoxyribose utilization in the mutant fitness data. There are many candidate genes in the genome, but none with that phenotype. We suspect that the conversion of D-glyceryl-CoA to D-glycerate is genetically redundant and therefore cannot be identified by assaying mutant strains with only one transposon insertion. The glycerate kinase of *P. simiae* (PS417_13970) is 51% identical to glycerate kinase 1 from *Escherichia coli (garK)*, which forms 2-phosphoglycerate (Bartsch et al. 2008;Zelcbuch et al. 2015). 2-phosphoglycerate is an intermediate in glycolysis and thus links the proposed pathway to central metabolism.

In the preceding analysis, we proposed reactions for all of the putative enzymes with strong negative phenotypes on deoxyribose (relative to other conditions), except for the acetyl-CoA C-acetyltransferase (PS417_10515). That enzyme converts acetoacetate (which is produced by the β-keto acid cleavage enzyme) back to acetyl-CoA, but acetoacetate must first be activated to acetoacetyl-CoA. Acetoacetate might be activated by a CoA-transferase (PS417_10525:PS417_10520) or a CoA-synthetase (there are several candidates in the genome). None of these genes are important for the utilization of deoxyribose, so we propose that this activity is genetically redundant. Overall, the proposed pathway accounts for all of the putative enzymes that are specifically important for deoxyribose utilization in *P. simiae*.

### Oxidation of deoxyribose to ketodeoxyribonate *in vitro*

We tested the activity of the deoxyribose dehydrogenase and the deoxyribonate dehydrogenase *in vitro*. (For details, see Appendix 1.) First, we overexpressed the *iorA*- and *iorB*-like subunits of deoxyribose dehydrogenase (PS417_10890 and PS417_10885) in *E. coli* and tested the resulting cell lysate. We found that this preparation oxidized deoxyribose to deoxyribonate with phenazine methosulfate as the electron acceptor. The lactonase (PS417_07255) was not necessary: deoxyribonolactone may hydrolyze spontaneously in water; or it might be hydrolyzed by enzymes in the cell lysate; or the deoxyribose dehydrogenase may form deoxyribonate instead of deoxyribonolactone. Second, we purified the deoxyribonate dehydrogenase (PS417_07245) and showed that it oxidized deoxyribonate with NADH as the electron acceptor. Tandem mass spectrometry showed that the oxidized product was a ketodeoxyribonate, but we were not able to determine which position was oxidized. We also tested the two dehydrogenases on each others’ substrates (deoxyribose dehydrogenase on deoxyribonate or deoxyribonate dehydrogenase on deoxyribose) and did not observe any activity. Thus, biochemical assays confirmed the proposed activities of deoxyribose dehydrogenase and deoxyribonate dehydrogenase.

### Accessory genes for deoxyribose catabolism in *P. simiae*

Based on the proposed pathway, we can explain the roles of the other genes with specific phenotypes on deoxyribose (Figure 1B). First, the putative operon that includes the lactonase, the SDR deoxyribonate dehydrogenase, and the β-keto acid cleavage enzyme also includes two genes from the major facilitator superfamily of transporters (MFS) (PS417_07260 and PS417_07265). We propose that these two genes are required for the uptake of deoxyribonate. We suspect that deoxyribose is oxidized in the periplasm, because the *iorB*-like subunit has a potential signal for tat-dependent export (RRRFLA) near its N terminus, and the associated cytochrome c protein has a membrane anchor and may be exported. If deoxyribose is oxidized in the periplasm, then deoxyribonate would be the substrate of the transporter when either deoxyribose or deoxyribonate is the carbon source, which would explain why these transporter genes were important in both conditions. But we do not understand why two different transporter genes appear to be required. These transporter genes are at the end of the putative operon, so polar effects are not likely.

Second, a *gntR*-like transcription factor (PS417_07240) is present at the beginning of the deoxyribonate oxidation cluster. By comparing the sequences upstream of this cluster and upstream of similar clusters from other strains of *Pseudomonas* or from *Burkholderia graminis*, we identified the conserved palindrome GTGATCAC. This sequence occurs at −47 to −40 relative to the predicted start codon of PS417_07240 and at similar positions relative to its homologs. The same motif was identified among the binding sites of the transcriptional activator AkgR from *Rhodobacter sphaeroides* (Imam et al. 2015), which is 32% identical to PS417_07240. We propose that PS417_07240 regulates the expression of the cluster by binding to this motif.

Third, the transcription factor PS417_13975 (*sdaR*) is probably an activator for the expression of the glycerate kinase (PS417_13970), which is downstream of *sdaR*. Similarly, a close homolog of PS417_13975 from *P. fluorescens* Pf-5 (PFL_3379, 88% identical) is predicted in RegPrecise (Novichkov et al. 2013) to regulate a downstream glycerate kinase.

Finally, *mobA* (PS417_21490) is expected to be necessary for the attachment of a nucleotide to molybdenum cofactor to form a molybdenum dinucleotide cofactor. This cofactor is probably required for the activity of deoxyribose dehydrogenase, which is homologous to the *coxLS* subunits of a CO dehydrogenase that has a molybdenum-cytosine dinucleotide cofactor (Dobbek et al. 1999). While *mobA* is required for the activity of deoxyribose dehydrogenase, it is not important for oxidation of deoxyribonate (Figure 1B), which is expected because none of the other enzymes in the pathway use a molybdenum cofactor. Similarly, genes for the biosynthesis of molybdopterin (a precursor to molybdenum cofactor) are important for utilizing deoxyribose but not deoxyribonate (*moaABCE*: PS417_21395, PS417_08670, PS417_04865, PS417_04875). These genes’ phenotypes are not specific to deoxyribose (and hence are not shown in Figure 1B) because they are also important for fitness in other conditions such as inosine utilization and nitrate stress (Price et al. 2018).

### Induction of the oxidative pathway in *P. simiae*

Next we asked if the enzymes in the oxidative pathway are induced when *P. simiae* grows on deoxyribose or deoxyribonate. As shown in Figure 2, all of these proteins seem to be expressed more highly during growth on deoxyribose or deoxyribonate than during growth on glucose. (The *iorA*-like subunit of deoxyribose dehydrogenase and the associated cytochrome were not detected; they may be membrane associated.) The differences in expression levels were statistically significant for the lactonase, the deoxyribonate dehydrogenase, and acetyl-CoA C-acetyltransferase (false discovery rate < 5%, ANOVA test).

**Figure 2:**
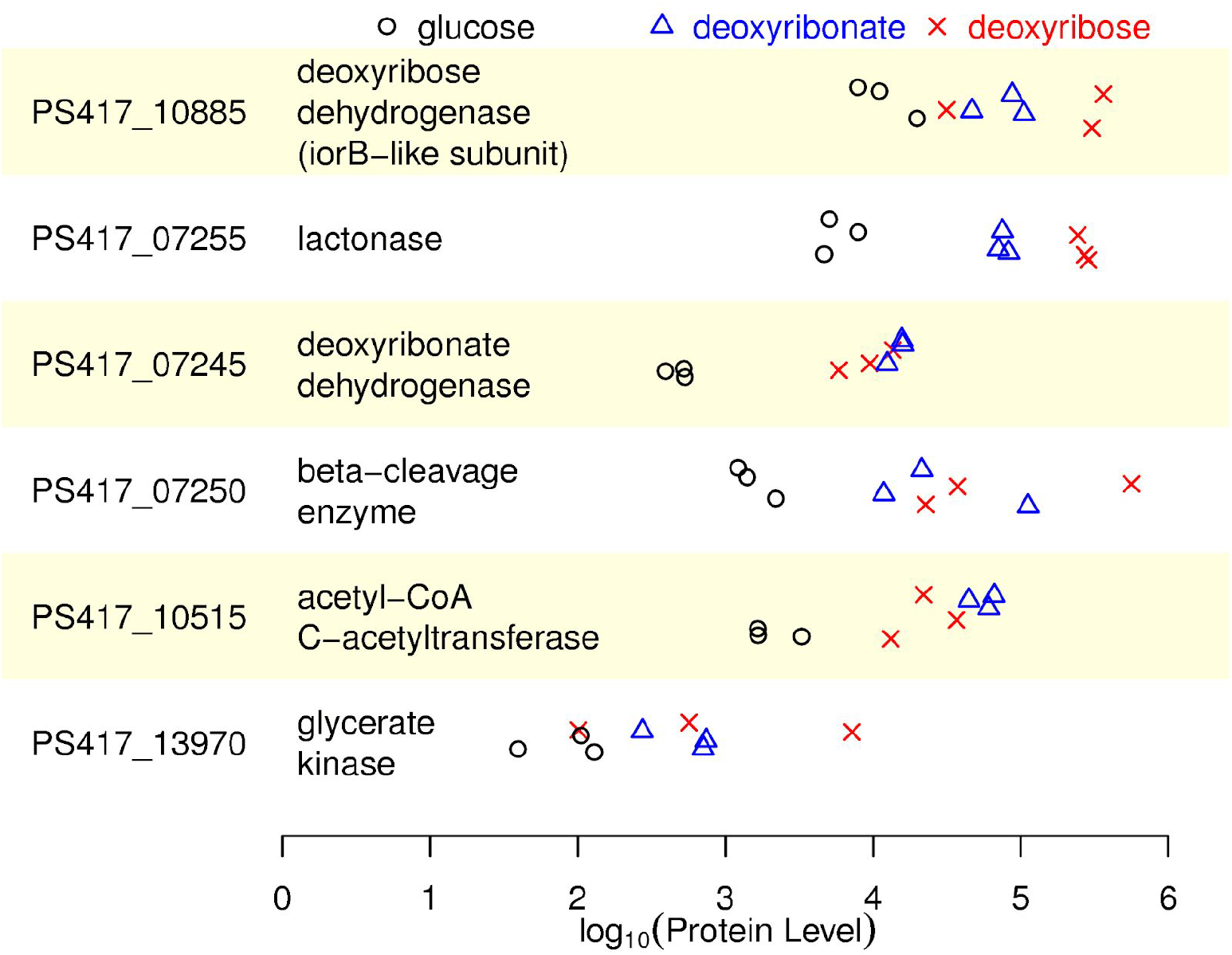
Levels of protein expression in *Pseudomonas simiae* WCS417. We show the normalized abundance of enzymes in the deoxyribose utilization pathway, as estimated from the integrated ion abundances of the detected peptide ions. There are three biological replicates of each condition and the *y* axis within each panel is random. Note the log scale for the *x* axis.

### Comparative genetics of the oxidative pathway

To determine if other bacteria use the oxidative pathway, we examined the genomic organization of the putative enzymes and their homologs. We focused on homologs from bacteria for which mutant libraries are available, and we obtained genome-wide mutant fitness data on deoxyribose, deoxyribonate, or deoxynucleosides for three additional bacteria: *Burkholderia phytofirmans* PsJN, *Paraburkholderia bryophila* 376MFSha3.1, and *Klebsiella michiganensis* M5al. This section gives an overview of the genes that were important for fitness in these conditions, the chromosomal organization of these genes, and the catabolic pathways that we propose.

Although these three additional bacteria, and *P. simiae*, grow on deoxyribonate by oxidative pathways, their other catabolic capabilities vary. Figure 3A shows the relationships between the proposed pathways for the catabolism of deoxyribose, deoxyribonate, or deoxynucleosides in the four bacteria based on our analysis of the genetics data. The full rationale for the assignment of each bacterium to a particular pathway is contained in the following sections.

**Figure 3:**
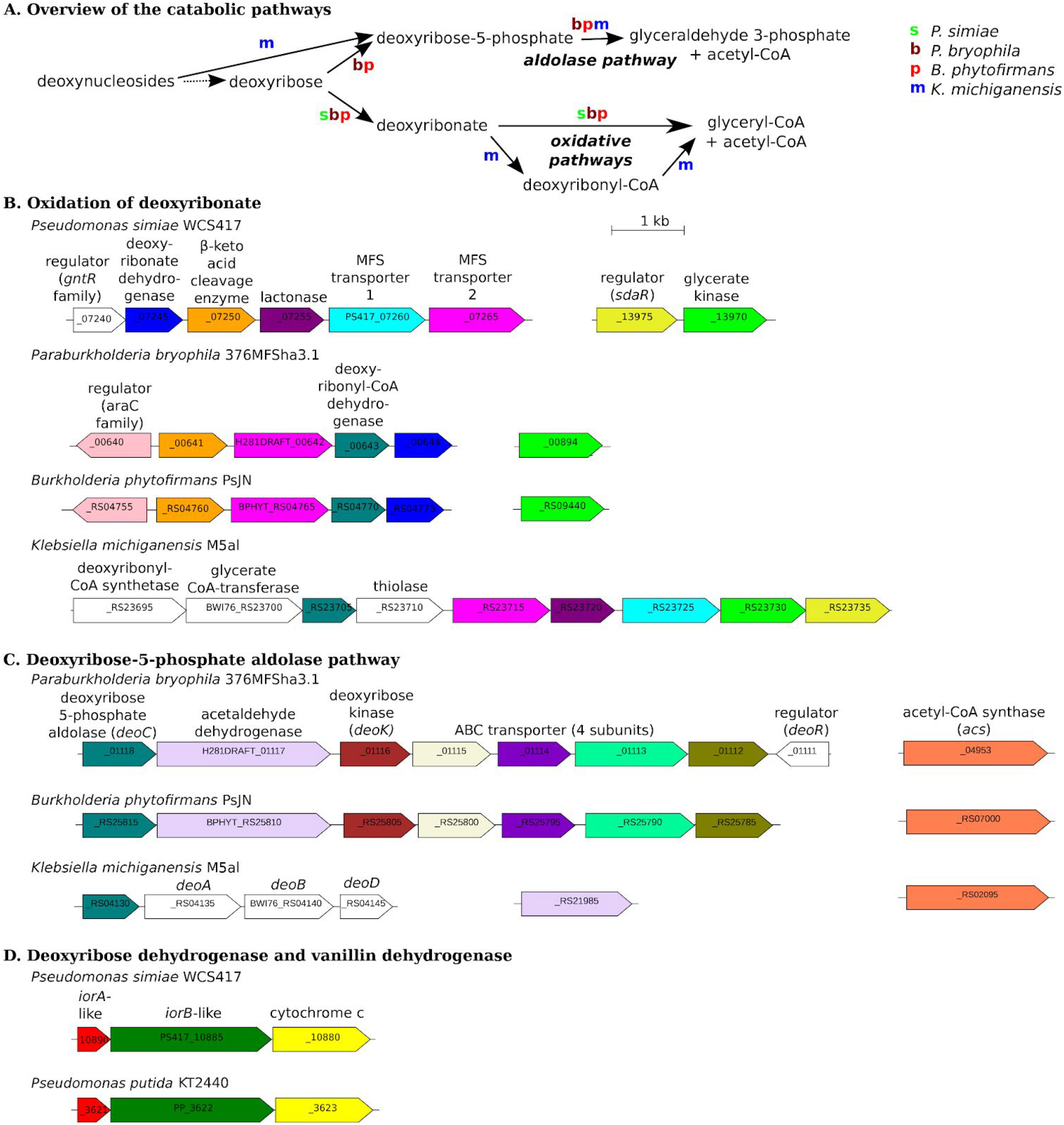
Genome organization of the deoxyribose utilization pathways. (A) Overview of the catabolism of deoxyribose and related compounds by 4 bacteria. For each transformation, we show which of the bacteria performs it. (B) Deoxyribonate oxidation in four bacteria. (C) The deoxyribose-phosphate aldolase pathway in three bacteria. (D) Deoxyribose dehydrogenase from *P. simiae* and vanillin dehydrogenase from *P. putida*. In panels B-D, adjacent genes are connected by horizontal lines, genes that are potential orthologs are the same color, and the scale is the same. All of the putative enzymes shown are important for the utilization of deoxyribose or related compounds except for the *iorAB*-like cluster in *P. putida*, the cleavage enzyme and deoxyribonyl-CoA dehydrogenase in *B. phytofirmans*, and we lack fitness data for *deoC* from *P. bryophila.*

First, the deoxyribonate oxidation genes are clustered together, with some variation, in diverse bacteria (Figure 3B). The clusters from two related bacteria, *Paraburkholderia bryophila* 376MFSha3.1 and *Burkholderia phytofirmans* PsJN, contain deoxyribonate dehydrogenase and the β-keto acid cleavage enzyme, and these genes were important for deoxyribonate utilization. The genomes of both of these bacteria also encode the kinase/aldolase pathway for deoxyribose utilization (Figure 3C). Genes from both pathways were important for deoxyribose utilization, so we propose that these bacteria consume deoxyribose by the kinase/aldolase and oxidative pathways in parallel.

Second, *Klebsiella michiganensis* M5al has a gene cluster that shares some genes with the deoxyribonate oxidation cluster, but *K. michiganensis* does not encode deoxyribonate dehydrogenase or the β-keto acid cleavage enzyme (Figure 3B). Nevertheless, this gene cluster was important for deoxyribonate utilization by *K. michiganensis*. We propose that *K. michiganensis* consumes deoxyribonate by another oxidative pathway with acyl-CoA intermediates. Also, *K. michiganensis* does not grow on deoxyribose, but it can grow on deoxynucleosides via DeoABD, which can convert them to deoxyribose-5-phosphate, and deoxyribose 5-phosphate aldolase (Figure 3C).

Third, the deoxyribose dehydrogenase of *P. simiae* is part of a conserved gene cluster, but its biological role does not seem to be conserved. The conserved cluster includes *iorA* and *iorB-*like subunits and a putative cytochrome c (Figure 3D). (In contrast, *iorAB* of isoquinoline 1-oxidoreductase are more distantly related, and no gene for a cytochrome c was found near *iorB* (Lehmann et al. 1995) or near its closest homologs in complete genomes (WP_083944220.1 or WP_113935277.1).) The deoxyribose dehydrogenase-like gene cluster is found in some other *Pseudomonas* species and in some genera of Proteobacteria such as *Bordetella* and *Agrobacterium*. Although the cluster is found in *Pseudomonas putida* KT2440, *P. putida* does not grow on deoxyribose or deoxyribonate, and its genome not contain the deoxyribonate oxidation genes. While studying the seemingly unrelated process of vanillin (4-hydroxy-3-methoxybenzaldehyde) catabolism by *P. putida*, we identified mutant phenotypes for this cluster. We propose that the cluster from *P. putida* (PP_3621:PP_3623) encodes a novel vanillin dehydrogenase.

We will now describe the genetic data for each of the bacteria in more detail.

### Oxidation of deoxyribose and deoxyribonate in *Paraburkholderia bryophila and* Burkholderia phytofirmans

*Paraburkholderia bryophila* 376MFSha3.1 and *Burkholderia phytofirmans* PsJN contain close homologs of the deoxyribonate dehydrogenase and the β-keto acid cleavage enzyme (50-60% identical to the proteins in *P. simiae*), but they do not contain close homologs of the deoxyribose dehydrogenase or the lactonase. In *P. bryophila,* the standard kinase/aldolase pathway was important for growth on deoxyribose, but not on deoxyribonate or glucose (Figure 4). Enzymes for the oxidative catabolism of deoxyribonate — the deoxyribonate dehydrogenase (H281DRAFT_00644) and the β-keto acid cleavage enzyme (H218DRAFT_00641) — were important for growth on both deoxyribose and deoxyribonate. Also important for growth on both substrates were glycerate kinase (H281DRAFT_00894) and genes for the conversion of acetoacetate to acetoacetyl-CoA to acetyl-CoA (H281DRAFT_04495 and H281DRAFT_00852). These genes were not important for glucose utilization. These data strongly suggest that the oxidative pathway in *P. bryophila* is the same as in *P. simiae.*

**Figure 4:**
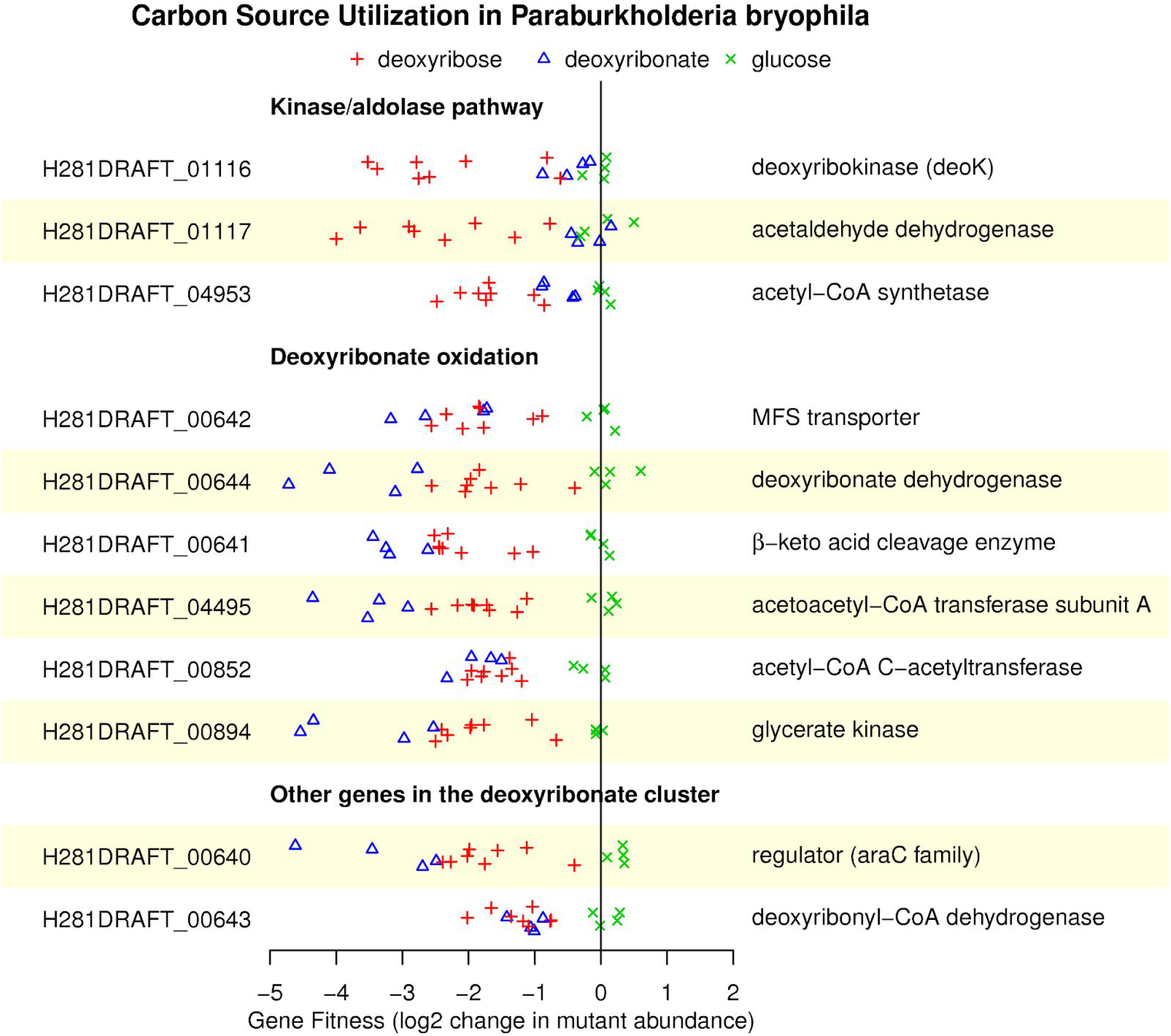
Mutant fitness data for the kinase/aldolase and oxidative pathways in *Paraburkholderia bryophila* 376MFSha3.1. Fitness values for the deoxyribose 5-phosphate aldolase (*deoC*) and for subunit B of acetoacetyl-CoA transferase were not estimated due to inadequate coverage. The data is plotted as in Figure 1B.

For most of the genes in the oxidative pathway, the mutant phenotypes were stronger (the fitness values were more negative) during growth on deoxyribonate than on deoxyribose (Figure 4). It appears that *P. bryophila* uses the oxidative pathway to grow on deoxyribonate, but deoxyribose is metabolized by the oxidative pathway in parallel with the kinase/aldolase pathway.

We did not identify any candidates for deoxyribose dehydrogenase in the fitness data from *P. bryophila*. This activity may be genetically redundant or non-specific, or the kinase/aldolase pathway may suffice for efficient growth on deoxyribose.

In the genome of *P. bryophila*, the oxidative enzymes are clustered with a transporter (H281DRAFT_00642), a regulator of the *araC* family (H281DRAFT_00640), and a putative β-hydroxyacyl-CoA dehydrogenase (H281DRAFT_00643) (Figure 3B). These genes are also important for the utilization of both deoxyribose and deoxyribonate, although mutants in the β-hydroxyacyl-CoA dehydrogenase have a milder phenotype (Figure 4). Based on the phenotype of a homolog of this protein in *K. michiganensis* (see below), we suspect that the β-hydroxyacyl-CoA dehydrogenase acts on deoxyribonyl-CoA. So, we speculate that *P. bryophila* may use a third parallel pathway for deoxyribose catabolism.

When we performed fitness assays in *B. phytofirmans*, we found similar results to *P. bryophila* for deoxyribonate utilization (Figure 5). In particular, deoxyribonate dehydrogenase (BPHYT_RS04775), the β-keto acid cleavage enzyme (BPHYT_RS04760), and glycerate kinase (BPHYT_RS09440) are all important during growth on deoxyribonate, although mutants of the β-keto acid cleavage enzyme have a milder phenotype. One unexplained result is that the acetyl-CoA C-acetyltransferase (BPHYT_RS09150) is detrimental to fitness on deoxyribonate (mutants in this gene increased more than 4-fold in abundance relative to other mutants in the pooled assay). During growth of *B. phytofirmans* on deoxyribose, we observe strong phenotypes for the enzymes in the kinase/aldolase pathway (Figure 5). Genes in the oxidative pathway are somewhat important for fitness during growth on deoxyribose at 10 mM (but not at 5 mM). Although these defects are modest, the defects are larger during growth on 10 mM deoxyribose than in most of 80 other conditions that we previously studied (data from (Price et al. 2018); see Figure 5).

**Figure 5:**
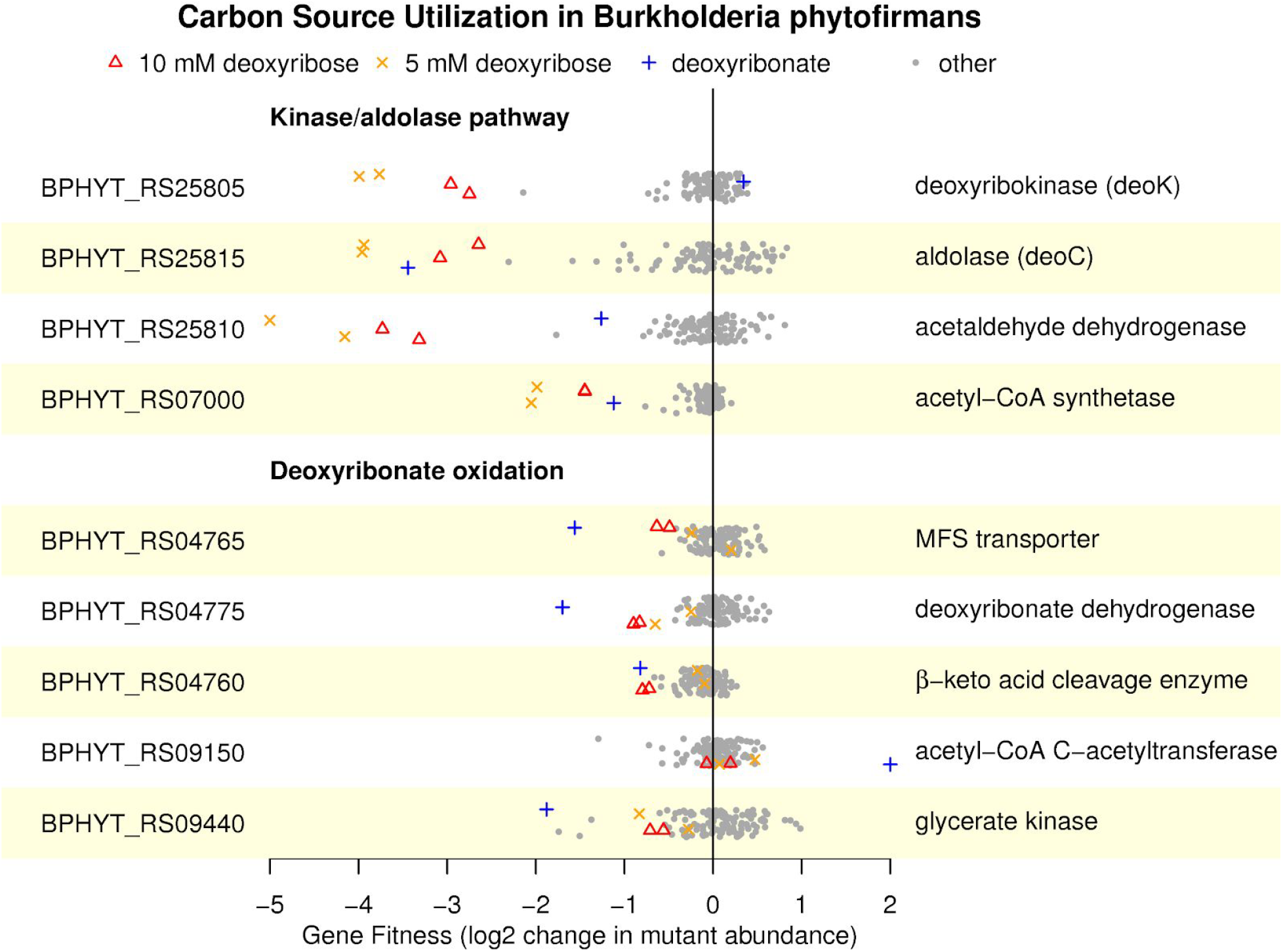
Mutant fitness data for the aldolase and oxidative pathways in *Burkholderia phytofirmans* PsJN. Fitness values outside of the plot’s range are shown at −5 or +2. The data is plotted as in Figure 1B.

Overall, we conclude that both *Paraburkholderia bryophila* and *Burkholderia phytofirmans* catabolize deoxyribonate by the same oxidative pathway that *P. simiae* uses. Furthermore, the oxidative pathway contributes to deoxyribose utilization in *P. bryophila* and probably in *B. phytofirmans* as well. However, in contrast to *P. simiae*, both *P. bryophila* and *B. phytofirmans* also use the aldolase-kinase pathway for deoxyribose catabolism.

### Oxidation of deoxyribonate in *Klebsiella michiganensis*

As mentioned above, the genome of *Klebsiella michiganensis* contains a gene cluster that shares some genes with the deoxyribonate oxidation cluster of *P. simiae* (Figure 3B). In *K. michiganensis*, the cluster is important for the utilization of deoxyribonate but not for the utilization of deoxynucleosides or deoxynucleotides (Figure 6) or in any of 92 other conditions that we previously profiled (Price et al. 2018). The cluster includes a putative CoA-synthetase (BWI76_RS23695), a β-hydroxyacyl-CoA dehydrogenase (BWI76_RS23705), a thiolase (BWI76_RS23710), a CoA-transferase (BWI76_23700), and glycerate kinase (BWI76_RS23730). We propose that the CoA-synthetase converts deoxyribonate to deoxyribonyl-CoA, the dehydrogenase oxidizes this to 3-ketodeoxyribonyl-CoA, and the thiolase cleaves this to glyceryl-CoA and acetyl-CoA (Figure S2). These transformations are analogous to the oxidative pathway of the other three bacteria, but with CoA bound intermediates. In all four bacteria, deoxyribonate is oxidized and cleaved to yield glyceryl-CoA and acetyl-CoA, but the point at which the CoA thioesterification occurs varies. In *K. michiganensis*, the cluster also includes a CoA-transferase that may convert the glyceryl-CoA to glycerate and a glycerate kinase that forms 2-phosphoglycerate, which feeds into glycolysis (as in the other bacteria).

**Figure 6:**
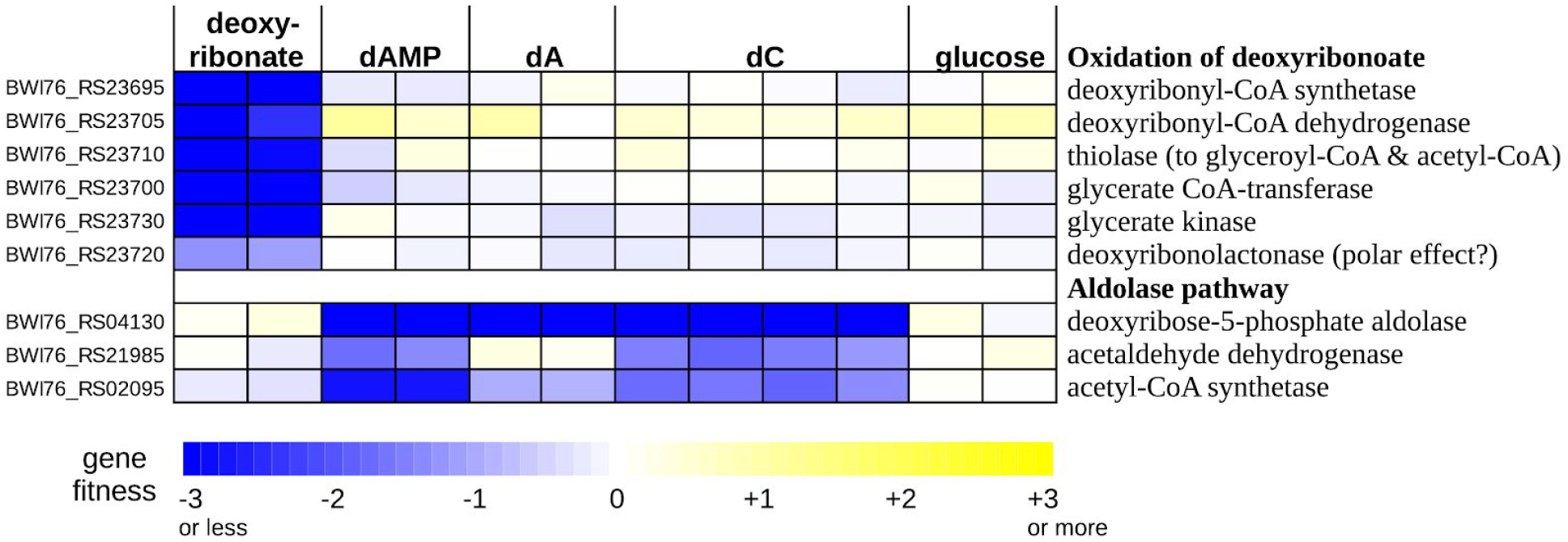
Mutant fitness data for the utilization of deoxyribonate and various deoxynucleotides/deoxynucleosides in *Klebsiella michiganensis* M5al. dAMP is 2’-deoxyadenosine 5’-phosphate, dA is 2’-deoxyadenosine, and dC is 2’-deoxycytidine.

The deoxyribonate utilization cluster of *K. michiganensis* also includes a close homolog (70% amino acid identity) of the putative deoxyribonolactonase from *P. simiae*. This gene (BWI76_RS23720) was also somewhat important for deoxyribonate utilization (fitness = −1.3 and −1.1). It is possible that the latter phenotype is due to polar effects that inhibit the expression of downstream genes (which include glycerate kinase). Or, because deoxyribonolactone and deoxyribonate interconvert in solution, the lactonase’s activity may be useful. We also observed a similar mild phenotype for the lactonase during deoxyribonate utilization in *P. simiae* (Figure 1B).

Although the genome of *K. michiganesis* contains a deoxyribose-phosphate aldolase (BWI76_RS04130), it does not seem to encode deoxyribose kinase. This might explain why it did not grow with deoxyribose as the carbon source. We did obtain fitness data with three deoxynucleotides or deoxynucleosides as carbon sources: 2’-deoxyadenosine 5’-phosphate, 2’-deoxyadenosine, and 2’-deoxycytidine (Figure 6). These data confirmed that *K. michiganensis* uses the aldolase pathway to break down deoxynucleosides. Deoxynucleosides are probably converted to deoxyribose 5-phosphate by the combination of a phosphorylase (either DeoA or DeoD; BWI76_RS04135 or BWI76_RS04145) which forms deoxyribose 1-phosphate and a phosphopentomutase (DeoB, BWI76_RS04140) which converts it to deoxyribose 5-phosphate. *DeoABD* were important for growth on deoxynucleotides and deoxynucleosides (all fitness values were under −1 except for *deoD* on 2’-deoxycytidine). Based on these data, we propose that *K. michiganensis* uses an oxidative pathway with acyl-CoA intermediates to metabolize deoxyribonate, it uses the aldolase pathway to metabolize deoxynucleosides, and it cannot metabolize deoxyribose.

### Deoxyribose dehydrogenase is closely related to a novel vanillin dehydrogenase from *Pseudomonas putida* KT2440

We found that all three genes from the deoxyribose dehydrogenase-like cluster of *Pseudomonas putida* KT2440 (PP_3621:PP_3623) were specifically important for growth on 5 mM vanillin (all fitness values ≤ −2 in two replicate experiments). These genes were not important for growth with other aromatic compounds as carbon sources (vanillic acid, benzoic acid, ferulic acid, or p-coumaric acid; Figure S3).

Because PP_3621:PP_3623 are important for growth on vanillin but not on vanillic acid (which is the oxidized form), we propose that it is the major vanillin dehydrogenase in *P. putida*. Although another vanillin dehydrogenase from *P. putida* (*vdh* or PP_3357) has been studied biochemically, *vdh* is not required for vanillin utilization (Simon et al. 2014). (Our data instead suggest a role for *vdh* as a p-hydroxybenzaldehyde dehydrogenase during the catabolism of p-coumaric acid (Figure S3).) Vanillin utilization by *P. putida* requires the uptake of molybdate (Graf and Altenbuchner 2014), which we can now explain because molybdate is a precursor for the molybdenum cofactor of PP_3621:PP_3623. The vanillin dehydrogenase is expected to produce vanillate, which is demethylated (by PP_3736:PP_3737) to protocatechuic acid, which feeds into the ortho-cleavage pathway that begins with protocatechuate 3,4-dioxygenase (PP_4655:PP_4656). The demethylase and dioxygenase genes are also specifically important during growth on vanillin (all fitness values under −2; Figure S3).

The genome of *P. putida* does not contain the deoxyribonate oxidation genes and *P. putida* does not grow with either deoxyribose or deoxyribonate as the sole carbon source. Conversely, none of the four other bacteria that we studied grew with vanillin as the sole carbon source, and their genomes do not encode orthologs of the vanillin O-demethylase. But all three components of *P. simiae*’s deoxyribose dehydrogenase are very similar to the components of *P. putida*’s vanillin dehydrogenase (84-92% amino acid identity). We speculate that both enzymes are active on a broader range of substrates.

## Discussion

### Oxidation and cleavage of deoxyribose and deoxyribonate

Using a genetic approach, we identified pathways for the oxidation of deoxyribose and/or deoxyribonate in four genera of bacteria. Although we have biochemical evidence for the conversion of deoxyribose to a ketodeoxyribonate in *Pseudomonas simiae,* we do not have direct evidence for the cleavage to D-glyceryl-CoA and acetyl-CoA. But the importance of glycerate kinase for deoxyribonate utilization by four different bacteria is difficult to explain otherwise. Also, for the three bacteria that use a β-keto cleavage enzyme during growth on deoxyribonate, experimental data from a homologous enzyme is consistent with our proposal. Specifically, *in vitro* assays of the β-keto cleavage enzyme BKACE_178 (UniProt A6X2V8), which is 61% identical to PS417_07250, found that it had some activity with 5-hydroxy-β-ketohexanoate, β-ketopentanoate, β-ketoisocaproate, and β-ketohexanoate as substrates (Bastard et al. 2014). These substrates are similar to 2-deoxy-3-keto-ribonate (which is 4,5-dihydroxy-β-ketopentanoate). Furthermore, the oxidative cluster in *Klebsiella michiganensis* contains glycerate kinase along with close homologs of two of the genes from the oxidative cluster of *P. simiae* (the lactonase and one of the MFS transporters; Figure 3B). This suggests that these gene clusters evolved to catabolize deoxyribonate via glycerate and 2-phosphoglycerate.

### What are the natural substrates of the oxidative pathways?

We identified deoxyribose dehydrogenase in just one of the four bacteria that grow on deoxyribonate. Furthermore, the natural role of the deoxyribose dehydrogenase of *P. simiae* remains uncertain. Its close similarity to the major vanillin dehydrogenase of *Pseudomonas putida* suggests that both enzymes might act on a broader range of substrates. And *P. simiae* grows more slowly on deoxyribose than on deoxyribonate (Figure S1), which suggests that deoxyribose might not be the natural substrate. Overall, it appears that deoxyribose utilization is not the main reason that these bacteria maintain a cluster for deoxyribonate oxidation.

We also wondered if 2-deoxyglucose might be a substrate for the oxidative pathway. A strain of *Pseudomonas* was reported to grow on 2-deoxyglucose by oxidizing it to 2-deoxy-3-ketogluconate (Eichhorn and Cynkin 1965). This transformation is chemically analogous to the conversion of 2-deoxyribose to 2-deoxy-3-ketoribonate. We tested the five bacteria considered here with 2.5-20 mM 2-deoxyglucose as a carbon source and found that none of them grew. Also, 2-deoxyglucose is not, as far as we know, a naturally occuring compound.

Instead, we propose that deoxyribonate is the natural substrate for the oxidative pathways. There are several plausible pathways by which deoxyribonate may form. First, non-specific oxidation of deoxyribose, which may occur in *Paraburkholderia bryophila* and in *Burkholderia phytofirmans*, could yield deoxyribonate. Although we did not identify the genes responsible for this non-specific activity, other sugar 1-dehydrogenase enzymes are known to be promiscuous, such as bifunctional L-arabinose/D-galactose 1-dehydrogenases (Price et al. 2018;Watanabe et al. 2006; Aro-Kärkkäinen et al. 2014). Also, because the deoxyribose dehydrogenase from *P. simiae* probably acts on substrates in the periplasm, some bacteria might obtain energy (but not carbon) by oxidizing deoxyribose in the periplasm without metabolizing it further. Deoxyribose in DNA is abundant, so even a low rate of non-specific oxidation could select for deoxyribonate catabolism. (All four of the deoxyribonate-catabolizing bacteria in this study are associated with plant roots. DNA is estimated to be ~3% of the dry weight of a typical plant cell (Landenmark et al. 2015), which implies that deoxyribose in DNA is around 1% of biomass.) Second, deoxyribonate or deoxyribonolactone could be released after oxidative damage of DNA (Kappen and Goldberg 1989). Third, deoxyribonate is proposed to form spontaneously from 2-keto-3-deoxy-D-gluconate (which is an intermediate in hexuronate metabolism and in variants of the Entner-Doudoroff pathway) and hydrogen peroxide (Lerma-Ortiz et al. 2016). Although we do not know of any measurement of the concentration of deoxyribonate in plants or in soils, 2-deoxyribonate is present in human urine at about 40-fold less than the concentration of creatinine (Chamberlin and Sweeley 1987) or around 25 mg/day excreted by the typical adult.

## Materials and Methods

### Strains and growth media

*Pseudomonas simiae* WCS417 and *Paraburkholderia bryophila* 376MFSha3.1 were provided by Jeff Dangl (University of North Carolina). Pools of barcoded transposon mutant of *Pseudomonas simiae* WCS417, *Pseudomonas putida* KT2440*, Burkholderia phytofirmans* PsJN, and *Klebsiella michiganensis* M5al were described previously (Price et al. 2018;Rand et al. 2017). Each mutant has a transposon insertion that includes *kanR* (providing kanamycin resistance) as well as a 20-nucleotide barcode flanked by common priming sites.

Individual strains from the library of mutants for *P. simiae* were obtained from an arrayed collection (Cole et al. 2017). Briefly, the library was spread on LB (Luria-Bertani) + kanamycin agar plates, and colonies were picked and arrayed into 384-well plates containing LB with 7.5% glycerol and kanamycin and allowed to grow overnight. Next, 25 nL of each well was collected as part of a multiplexing strategy involving pooling of rows, columns, and plates, and these pools were subjected to PCR amplification and sequencing of the insertion barcodes. The location of each mutant was determined using an in-house script to identify the row, column, and plate pools sharing the same barcode. The insertion barcodes of the specific mutants used in this study were confirmed by Sanger sequencing of colonies recovered from glycerol stocks, using PCR with BarSeq_P1 and BarSeq_P2_noindex primers to amplify the region and BarSeq_P1_SangerSeq as the sequencing primer (Table 1). Furthermore, the presence of an insertion within the expected gene was confirmed by PCR amplification of the transposon junction using a mariner transposon-specific primer and a gene-specific primer (see the confirm primers in Table 1).

**Table 1:**
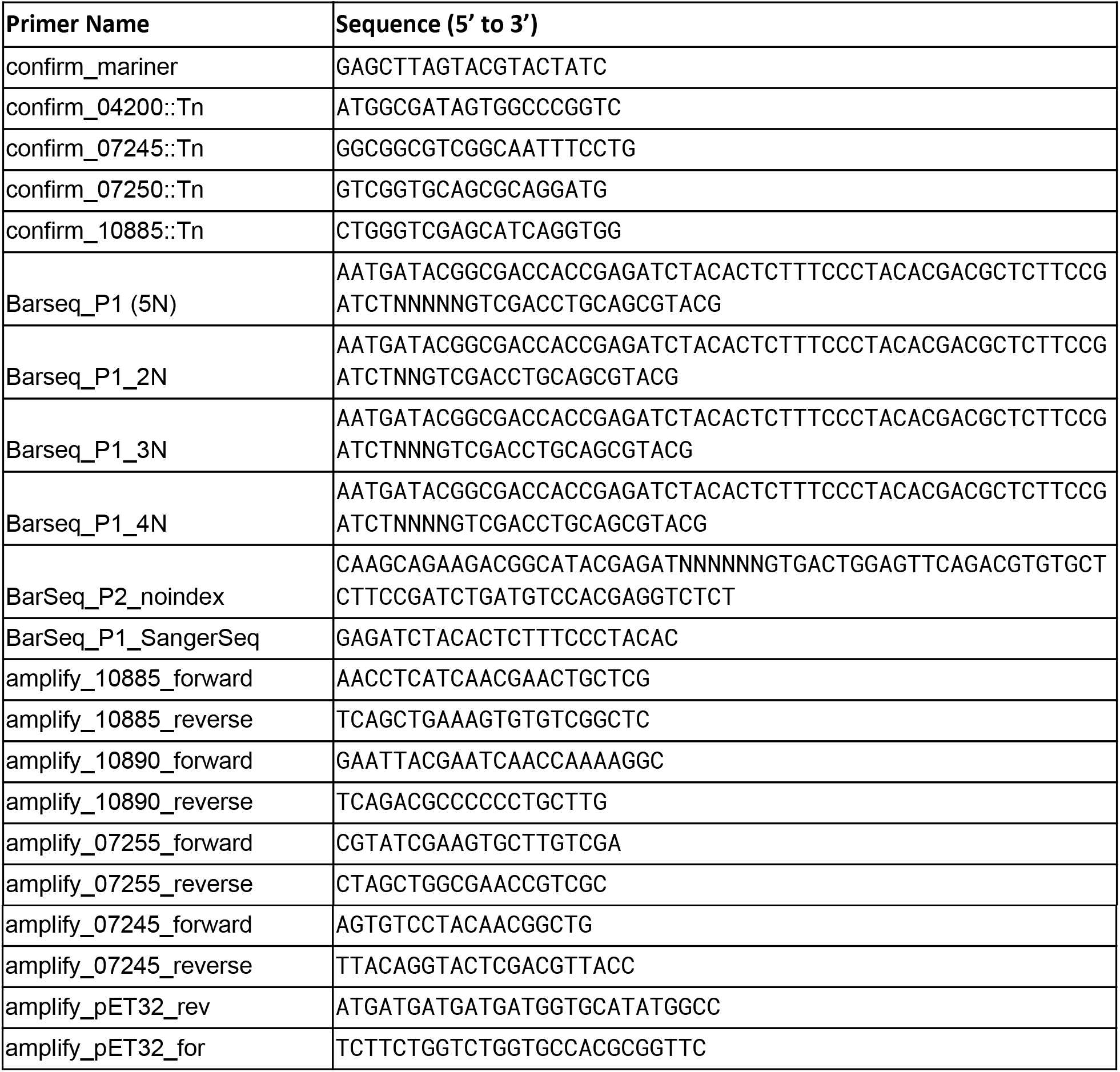
Primers used in this study.

A transposon mutant library of *Paraburkholderia bryophila* 376MFSha3.1 (Burk376_ML3) was constructed via conjugation with an *E. coli* donor strain (APA752) carrying the randomly barcoded mariner transposon delivery vector pKMW3, as described previously (Wetmore et al. 2015). Briefly, we conjugated a 1:1 ratio of mid-log phase *P. byrophila* to mid-log phase APA752 for 12 hours at 30°C on LB plates supplemented with diaminopimelic acid (DAP). The *E. coli* donor strain is derived from WM3064, a DAP auxotroph. After conjugation, we selected for mutants by plating on LB plates supplemented with 100 μg/mL kanamycin at 30°C. After pooling about 100,000 colonies, we sub-inoculated the cells into 50 mL of LB with 100 μg/mL kanamycin at a starting OD (optical density at 600 nm) of 0.2, grew the library to saturation at 30°C, and made multiple freezer stocks in glycerol. The mapping of transposon insertion locations and the identification of their associated DNA barcodes were performed as described previously (Wetmore et al. 2015). Burk376_ML3 contains 106,195 mapped strains with unique DNA barcodes and contains at least one central insertion (in 10-90% of the gene’s length) for 5,624 of 6,555 predicted protein-coding genes. Of 63,242 strains with insertions in the central parts of genes, 59,045 were at sufficient abundance in the recovered library and were used to compute gene fitness.

All bacteria were cultured in LB medium during recovery from glycerol stocks. For the recovery of mutant libraries or individual mutant strains, 50 μg/mL kanamycin was added to the LB. All mutant libraries except *K. michiganensis* were recovered until the cells reached mid-log phase (OD600 around 1.0). For *K. michiganensis*, we recovered the mutant library until the culture reached saturation (OD600 around 4.0). All bacterial growth described in this study was conducted at 30°C. For cloning work, LB was used as the rich medium. All chemicals for media preparations, mutant fitness assays, growth assays and biochemical reactions were purchased from Sigma unless otherwise mentioned. For deoxyribonate, we used the lithium salt.

For assaying fitness or growth on a specific carbon source, we generally used a defined medium with 5 − 20 mM of the carbon source as well as 30 mM PIPES sesquisodium salt, 0.25 g/L ammonium chloride, 0.1 g/L potassium chloride, 0.6 g/L sodium phosphate monobasic monohydrate, Wolfe’s vitamins, and Wolfe’s minerals (http://fit.genomics.lbl.gov). For some fitness assays of *P. putida*, we instead used a minimal MOPS medium (40 mM 3-(N-morpholino)propanesulfonic acid, 4 mM tricine, 1.32 mM potassium phosphate dibasic, 10 μM iron (II) sulfate heptahydrate, 9.5 mM ammonium chloride, 0.276 mM aluminum potassium sulfate dodecahydrate, 0.5 μM calcium chloride, 0.525 mM magnesium chloride hexahydrate, 50 mM sodium chloride, 3 nM ammonium heptamolybdate tetrahydrate, 0.4 uM boric acid, 30 nM cobalt chloride hexahydrate, 10 nM copper (II) sulfate pentahydrate, 80 nM manganese (II) chloride tetrahydrate, and 10 nM zinc sulfate heptahydrate).

For the growth assays with individual mutants of *P. simiae* (Figure S1), we washed the cells 3 times with defined media before starting the growth experiments (the cells were pre-grown in LB). We grew these individual strains in a 96 well microplate within a Tecan Sunrise instrument, with readings every 15 minutes, as described (Price et al. 2018).

### Genome-wide mutant fitness assays

For pooled fitness assays (RB-TnSeq), cells from the recovered mutant library were washed 3 times with defined media without a carbon source and then inoculated into the medium of choice at OD_600_ = 0.02 and grown until saturation in a transparent 24-well microplate plate in a Multitron shaker. Total genomic DNA was extracted and barcodes were PCR amplified as described previously (Wetmore et al. 2015), but with some modifications to the primers. For most of the experiments, the P1 primer included variable spacing between the Illumina adapter and the sequence that flanks the barcode (2-5 Ns instead of always 5 Ns; see Table 1). Barcodes were sequenced using an Illumina HiSeq 4000, which can lead to some bleed-through between samples (in our experience, 0.1% to 0.3% of reads from BarSeq; for the likely mechanism behind bleed-through, see (Sinha et al. 2017)). To eliminate this bleed-through, some experiments used a P1 primer that contained an additional sequence to verify that it came from the expected sample. (The P2 primer contains the tag that is used for demultiplexing by Illumina software; in our earlier design, the P1 primer was the same mix for all the samples.) Specifically, the new P1 primers contain the same sequence as BarSeq_P1 before the Ns, 1-4 Ns (which varies with the index), the reverse (not the reverse complement) of the 6-nt index sequence, and the sequence from BarSeq_P1 after the Ns. These 96 primer sequences are available with the analysis code (see metadata/barseq3.index2 at https://bitbucket.org/berkeleylab/feba).

### Analysis of the mutant fitness data

The fitness data was analyzed as described previously (Wetmore et al. 2015) using publically available scripts (https://bitbucket.org/berkeleylab/feba). Briefly, the fitness of a strain is the normalized log_2_ ratio of the number of reads for its barcode in the sample after growth versus the control sample (taken after the cells recover from the freezer). The fitness of a gene is the weighted average of the fitness of strains with insertions in the central 10-90% of the gene. For example, fitness values were computed for 4,414 of the 5,033 predicted non-essential proteins in *P. simiae*, and for the typical (median) protein, the fitness values of five independent mutant strains were averaged. For each experiment, the gene fitness values are normalized so that the mode of the distribution is at zero.

To assess the statistical significance of each fitness value, we used a *t*-like test statistic of the form fitness / sqrt(estimated variance) (Wetmore et al. 2015). For most of the bacteria, a gene was considered to have a specific phenotype in an experiment if fitness < −1, *t* < −5, 95% of fitness values are between −1 and 1, and |fitness| > 95th percentile(|fitness|) + 0.5, as described previously (Price et al. 2018). However, for *P. simiae*, 9/162 experiments (6%) are for deoxyribose or deoxyribonate, so the 95th percentile threshold for a specific phenotype was not appropriate. We used the 90th percentile instead.

In *P. simiae,* we found 19 genes with a specific phenotype in at least one of six deoxyribose experiments. We manually excluded 6 of these genes from Figure 1B because they did not have a consistent fitness defect. These genes were PS417_08675 or *moaA,* which is involved in molybdenum cofactor biosynthesis; PS417_23250 or *tsaA,* which is involved in tRNA modification; and four regulatory or signalling genes: PS417_00445, PS417_07615, PS417_13295, and PS417_22670.

### Sequence analysis

Protein sequences were analyzed by using PaperBLAST (Price and Arkin 2017) to search for characterized homologs; with the Fitness Browser (http://fit.genomics.lbl.gov; (Price et al. 2018)), which incorporates results from PFam (Finn et al. 2014), TIGRFam (Haft et al. 2013), KEGG (Kanehisa et al. 2016), and SEED/RAST (Overbeek et al. 2014); by using the Conserved Domain Database (Marchler-Bauer et al. 2015) to search for similarity to gene families; and by using PSORTb (Yu et al. 2010) to predict protein localization.

To identify a motif for the putative binding sites of the transcription factor PS417_07240, we first used the MicrobesOnline tree-browser (Dehal et al. 2010) to identify orthologs with conserved synteny. We identified similar proteins from *P. fluorescens* SBW25, *P. fluorescens* Pf-5, *P. syringae* B728a, and *B. graminis* C4D1M (44-99% amino acid identity) that were also upstream of an SDR dehydrogenase and a β-keto acid cleavage enzyme. We ran MEME (Bailey and Elkan 1994) on the 200 nucleotides upstream of these five genes.

### Cloning and protein purification

We used the expression vector pET32a for cloning and protein purification as described previously (Baran et al. 2013). We PCR amplified four genes from *P. simiae* WCS417 (PS417_10885, PS417_10890, PS417_07255, and PS417_07245) using the amplify primers listed in Table 1 and cloned the PCR products into pET32a using Gibson assembly (Gibson 2011). For PS417_07255, PS417_07245, and PS417_10890, we transformed the Gibson assembly products directly into the *E. coli* expression strain BL21(DE3). However, for PS417_10885 we first transformed into *E. coli* TOP10 to identify the correct plasmid. We then transformed this plasmid into the *E. coli* BL21(DE3) expression strain. Protein was purified using a cobalt affinity column (Clonetech TALON kit) and quantified using a Nanodrop spectrophotometer (Thermofisher). Purified PS417_07245 and PS417_07255 had the expected molecular weights (29 kDa and 39 kDa, respectively). Although we were not able to purify PS417_10885 or PS417_10890, protein gels of cell lysates from *E. coli* strains that expressed these proteins showed predominant bands at the expected molecular weights (80 kDa and 17 kDa, respectively).

### Enzymatic assays

Enzymatic assays were performed with purified enzymes (2-5 μM) except that clarified cell lysates of *E. coli* expressing PS417_10885 or PS417_10890 were mixed together to obtain deoxyribose dehydrogenase. Each enzymatic assay was done in 50 μL of 100 mM ammonium bicarbonate (NH_4_HCO_3_) buffer with three independent replicates, run for one hour at room temperature, and then diluted 1000-fold in ammonium bicarbonate buffer and frozen at −20°C. NADH oxidase was purchased from Nacalai USA (San Diego, CA).

### Liquid chromatography and mass spectrometry of end products

Samples were analyzed using a Thermo-Dionex UltiMate3000 RSLCnano liquid chromatograph that was connected in-line with an LTQ-Orbitrap-XL mass spectrometer equipped with a nanoelectrospray ionization (nanoESI) source (Thermo Fisher Scientific, Waltham, MA). The LC was equipped with a C18 analytical column (length: 150 mm; inner diameter: 0.075 mm; particle size: 3 μm; pore size: 100 Å; Thermo Acclaim®) and a 1-μL sample loop. Acetonitrile, formic acid (Optima grade, 99.9%, Fisher), and water purified to a resistivity of 18.2 MΩ⋅cm (at 25°C) using a Milli-Q Gradient ultrapure water purification system (Millipore, Billerica, MA) were used to prepare mobile phase solvents. Solvent A was 99.9% water/0.1% formic acid and solvent B was 99.9% acetonitrile/0.1% formic acid (v/v). The elution program consisted of isocratic flow at 1% B for 3 min, a linear gradient to 95% B over 1 min, isocratic flow at 95% B for 3 min, and isocratic flow at 1% B for 13 min, at a flow rate of 300 nL/min. Full-scan mass spectra were acquired in the positive ion mode over the range *m*/*z* = 70 to 1500 using the Orbitrap mass analyzer, in profile format, with a mass resolution setting of 60,000 (at *m/z* = 400, measured at full width at half-maximum peak height). For tandem mass spectrometry (MS/MS) analysis, precursor ions were fragmented using collision-induced dissociation (CID) under the following conditions: MS/MS spectra acquired using the linear ion trap, centroid format, isolation width: 2 m/z units, normalized collision energy: 35%, default charge state: 1+, activation Q: 0.25, and activation time: 30 ms. Data acquisition and analysis were performed using Xcalibur software (version 2.0.7, Thermo).

### Proteomics

*P. simiae* was inoculated at an initial OD_600_ of 0.04 into 1 mL of defined media that contained 10 mM of either glucose, deoxyribose or lithium deoxyribonate. At late log phase (OD_600_ = 0.4-0.7), cells were pelleted by centrifugation at 4,000 x g and resuspended in 100 mM NH4HCO3 pH 7.5 buffer (Sigma Aldrich, St. Louis, MO, USA). 50 μL of a 1 μg/μL sample of each cell suspension was mixed with 10 μL 100 mM NH_4_HCO_3_, pH 7.5 and 25 μL of a 0.2% aqueous solution of Rapigest SF (Waters, Milford, MA, USA) and the sample was placed in a block heater at 80°C for 15 minutes and vortexed briefly to lyse cells and solubilize proteins. 5 μL of a 1 μg/μL solution of Trypsin Gold Porcine Protease (Promega, Madison, WI, USA) was added and the sample was incubated overnight at 37°C for the tryptic digest of proteins. The next morning, 10 μL of trifluoroacetic acid (Sigma Aldrich) was added and the sample was incubated for 90 minutes at 37°C to hydrolyze the Rapigest SF. The sample was centrifuged at 21,130 x g for 30 minutes at 6°C, and the supernatant was transferred to an autosampler vial for mass spectrometry analysis.

Trypsin-digested proteins were analyzed using a Synapt G2-S*i* high-definition ion mobility mass spectrometer that was equipped with a nanoelectrospray ionization source and connected in line with an Acquity M-class ultra-performance liquid chromatography system (UPLC; Waters, Milford, MA). Data-independent, ion mobility-enabled mass spectra and tandem mass spectra (Plumb et al. 2006;Shliaha et al. 2013;Distler et al. 2014) were acquired in the positive ion mode. Data acquisition was controlled using MassLynx software (version 4.1). Tryptic peptide identification, relative protein quantification using a label-free approach (Neilson et al. 2011;Nahnsen et al. 2013) and calculation of statistical analysis of variance (ANOVA) p-values were performed using Progenesis QI for Proteomics software (version 4.0, Waters). The abundance of 1,277 proteins was quantified.

### Data availability

The fitness data is available from the Fitness Browser (http://fit.genomics.lbl.gov). The fitness data and the proteomics data are archived at figshare (https://doi.org/10.6084/m9.figshare.7304159.v1).

## Appendix 1 Oxidation of deoxyribose to a ketodeoxyribonate in vitro

To test the activity of deoxyribose dehydrogenase from *P. simiae in vitro*, we recombinantly expressed the *iorA-*like and *iorB*-like components (PS417_10890 and PS147_10885) with polyhistidine (6X His) tags in *E. coli*. We tried to use a cobalt affinity column to purify each subunit, but did not succeed. Instead, we combined cell lysates from two strains of *E. coli* that overexpressed each subunit.

We incubated these cell lysates with 1 mM deoxyribose and with 100 μM phenazine methosulfate as the electron acceptor. The *iorAB*-like dehydrogenase converted deoxyribose to a compound with the same mass-to-charge ratio (*m/z*) as deoxyribonate (Figure S4). Tandem mass spectrometry confirmed that this product had a fragmentation pattern very similar to that of a 2-deoxy-D-ribonate standard (Figure S5). In contrast, addition of an equivalent amount of cell lysate from *E. coli* with an empty vector yielded only a small amount of deoxyribonate (less than a tenth as much; Figure S4). This shows that the deoxyribose dehydrogenase activity was primarily due to the *iorAB*-like dehydrogenase from *P. simiae* and not to an enzyme that is native to *E. coli.* No deoxyribonate formed if no lysate was added, so the conversion does not occur spontaneously in solution. Using polyhistidine tags, we were able to purify the lactonase (PS417_07255). The conversion of deoxyribose to deoxyribonate occurred with or without the added lactonase, perhaps because deoxyribonolactone in water can spontaneously hydrolyze to deoxyribonate or because lactonases are present in the cell lysate. In both reactions, and in the starting sample of deoxyribose, we detected a small amount of a compound with *m/z* = 155.0 that could be deoxyribonolactone, although the abundance of this ion was too low for tandem mass spectrometry.

We also purified the deoxyribonate dehydrogenase (PS417_07245) using a polyhistidine tag and a cobalt affinity column. We incubated the purified enzyme with 1 mM deoxyribonate and with 1 mM NAD^+^ as the electron acceptor. We observed a product with the molecular weight of 2-deoxy-3-keto-D-ribonate (a putative sodium adduct with *m/z* = 171.02). The putative ketodeoxyribonate was not detected in the absence of the deoxyribonate dehydrogenase or when either deoxyribose or deoxyribonate was incubated with the *iorAB*-like dehydrogenase. Tandem mass spectrometry showed the loss of CO_2_ or H_2_O, as expected for a sugar acid (Figure S6). The amount of product was small and difficult to quantify (perhaps 3-4% of the ion intensity of the input deoxyribonate), but in the absence of the enzyme, the product was below the limit of detection. We also tested the deoxyribonate dehydrogenase for activity with deoxyribose as the substrate, but we did not observe any formation of deoxyribonate or ketodeoxyribonate.

Because the deoxyribonate dehydrogenase reaction is similar to the 2-deoxygluconate 3-dehydrogenase reaction, which is thermodynamically unfavorable (Eichhorn and Cynkin 1965), we tried to reoxidize the NADH in order to drive the reaction forward. Specifically, we incorporated 0.002 units/μL of NADH oxidase, which converts NADH + O_2_ to NAD^+^ + H_2_O_2_. This change increased the yield of the putative ketodeoxyribonate product to 12-14%. The retention time of the product in the HPLC was identical to that of deoxyribonate, so we did not try to separate them.

## Acknowledgements

We thank Mitchell Thompson for growing the *P. putida* mutant library on several carbon sources.

## Supplementary Figures

**Figure S1:**
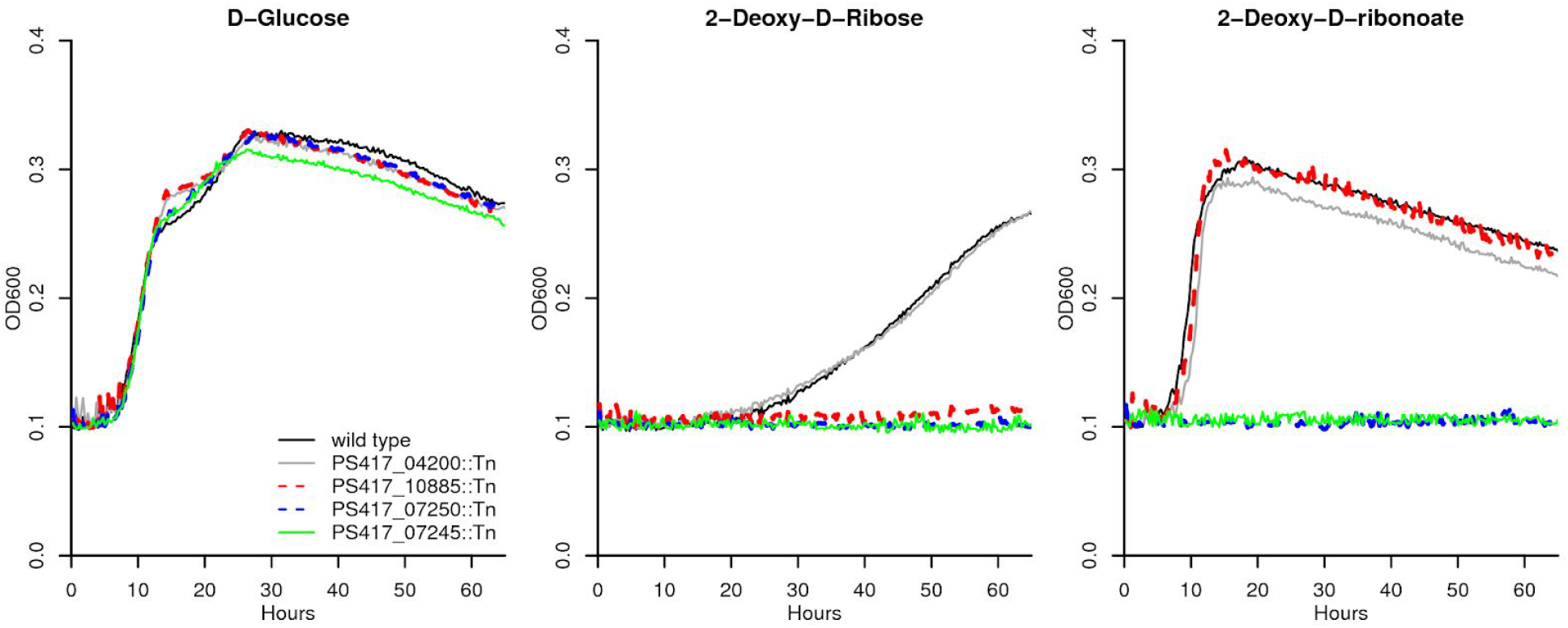
Growth curves for individual transposon mutants. We compared the growth of wild-type *P. simiae* WCS417 and of individual transposon mutant strains in defined media with three carbon sources: D-glucose, deoxyribose, and deoxyribonate. The mutant of PS417_04200 (2-ketoglutaric semialdehyde dehydrogenase) is included as a control. Each curve is the median of six replicates. Each carbon source was provided at 10 mM.

**Figure S2:**
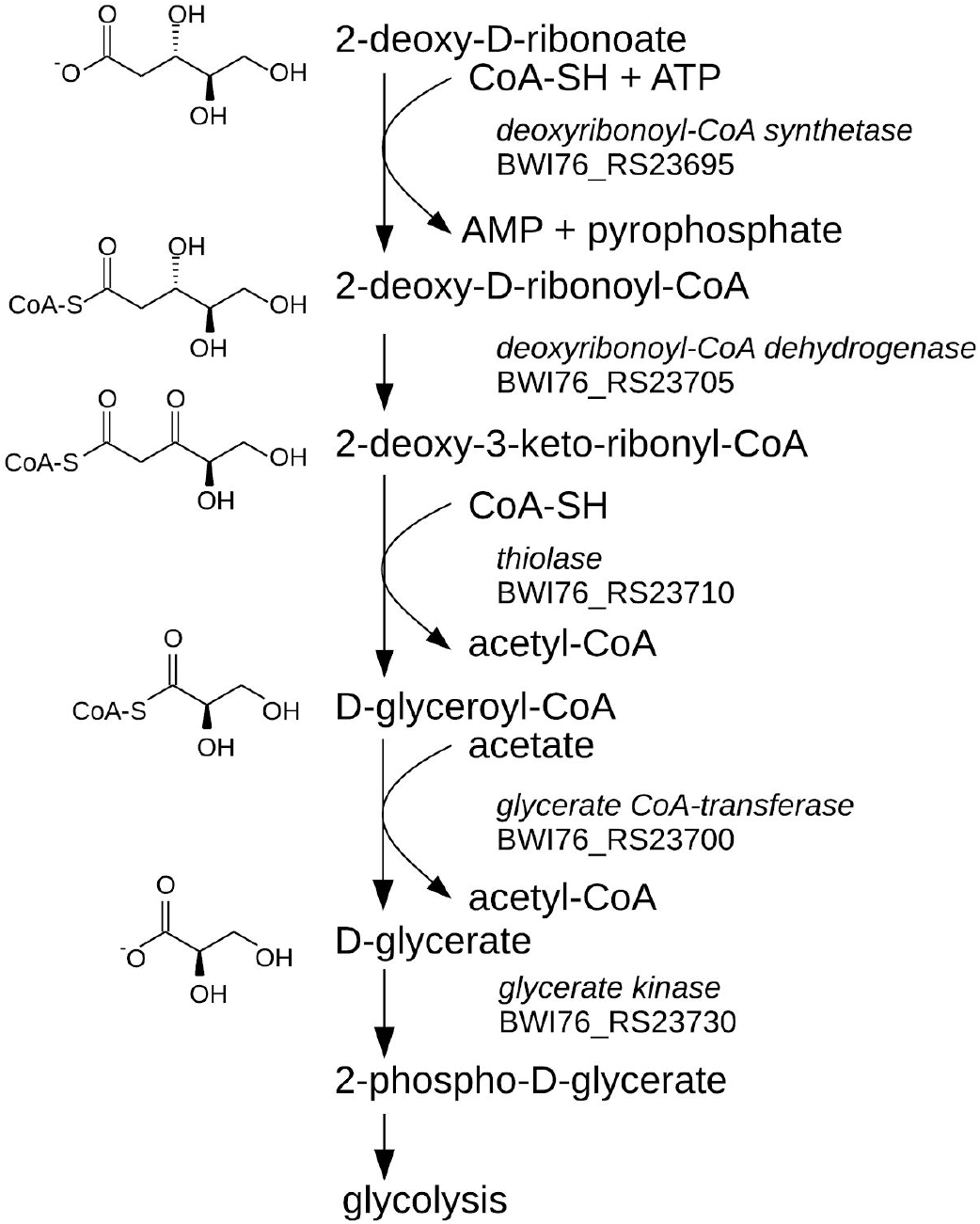
The proposed pathway of deoxyribonate oxidation in *Klebsiella michiganensis* M5al. We identified the closest characterized homolog of each putative enzyme using PaperBLAST (Price and Arkin 2017). The CoA-synthetase is 30% identical to the long-chain fatty-acid CoA-synthetase of *B. subtilis* (*lcfB* or O07610). The dehydrogenase is 45% identical to a 3-oxoacyl-[acyl-carrier-protein] reductase (*fabG* or P0A0H9), which carries out a reaction in the opposite (reductive) direction than shown above, and with the CoA attached to acyl carrier protein. However an ACP substrate is normally part of biosynthesis, which is difficult to reconcile with this gene’s mutant phenotype or its presence in the deoxyribonate utilization cluster. The thiolase is 47-50% identical to thiolases that act on acetoacetyl-CoA (see P45359 in Swiss-Prot or see PfGW456L13_2411 in the Fitness Browser). The CoA-transferase is 42% identical to a propionate CoA-transferase (Q9L3F7) that uses propionyl-CoA and acetate as substrates, so we show acetate as the CoA acceptor, but it might use another acceptor such as succinate.

**Figure S3:**
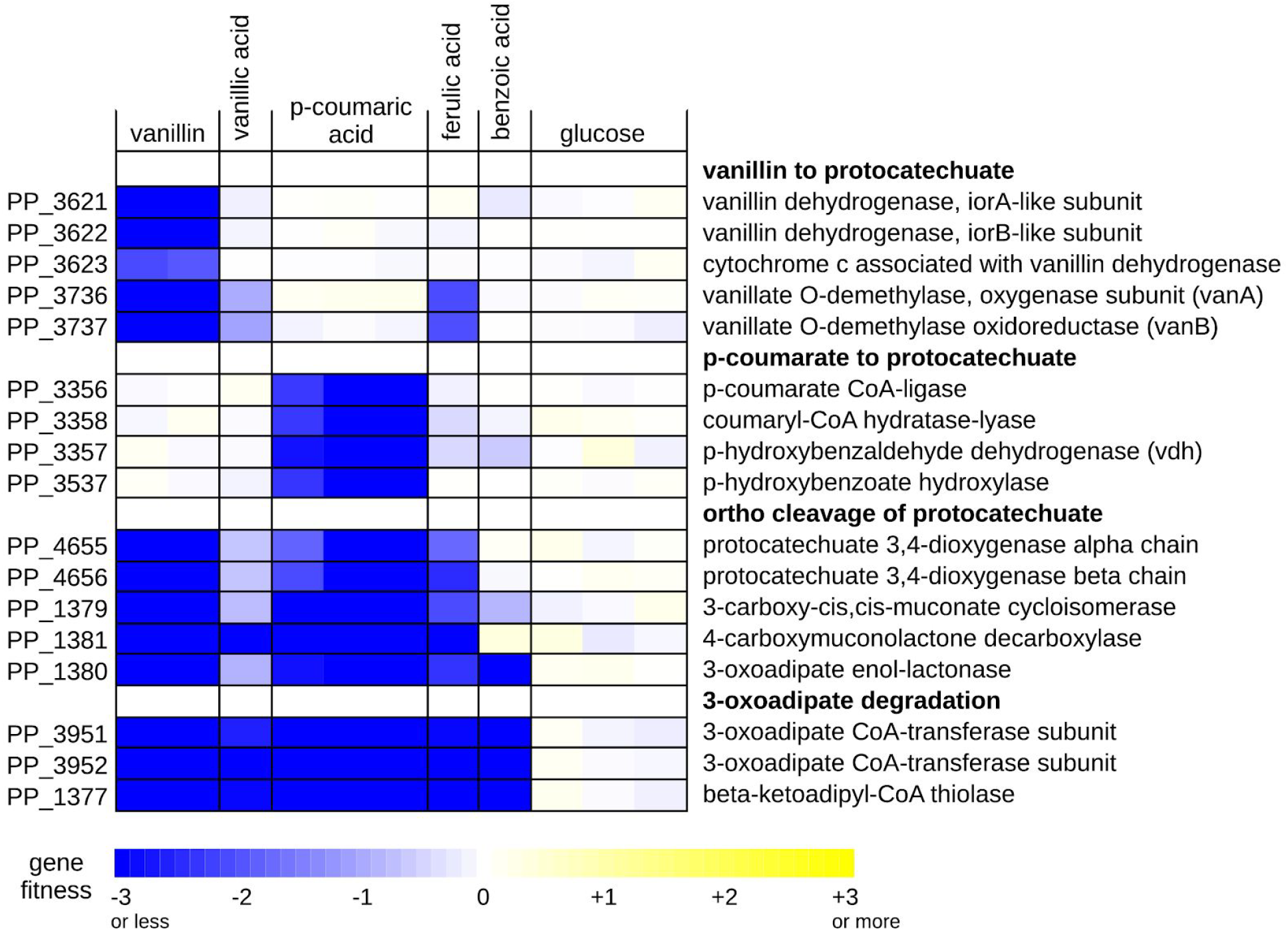
Fitness data for aromatic compound degradation by *Pseudomonas putida* KT2440. Vanillin (4-hydroxy-3-methoxybenzaldehyde) is oxidized to vanillate, demethylated to protocatechuate (3,4-dihydroxybenzoate), and catabolized via the ortho-cleavage pathway and 3-oxoadipate to yield acetyl-CoA and succinyl-CoA. p-coumarate (3-(4-hydroxyphenyl)-2-propenoic acid; also known as 4-hydroxycinnamic acid) appears to be activated to p-coumaryl-CoA, hydrated and cleaved to p-hydroxybenzaldehyde, oxidized to p-hydroxybenzoate, and hydroxylated to protocatechuate. Since mutants of *vdh* (PP_3357) are deficient in growth on p-coumaric acid, its primary substrate is probably p-hydroxybenzaldehyde, not vanillin. (The right-most glucose experiment, shown here as a control, is from (Thompson et al. 2018).)

**Figure S4:**
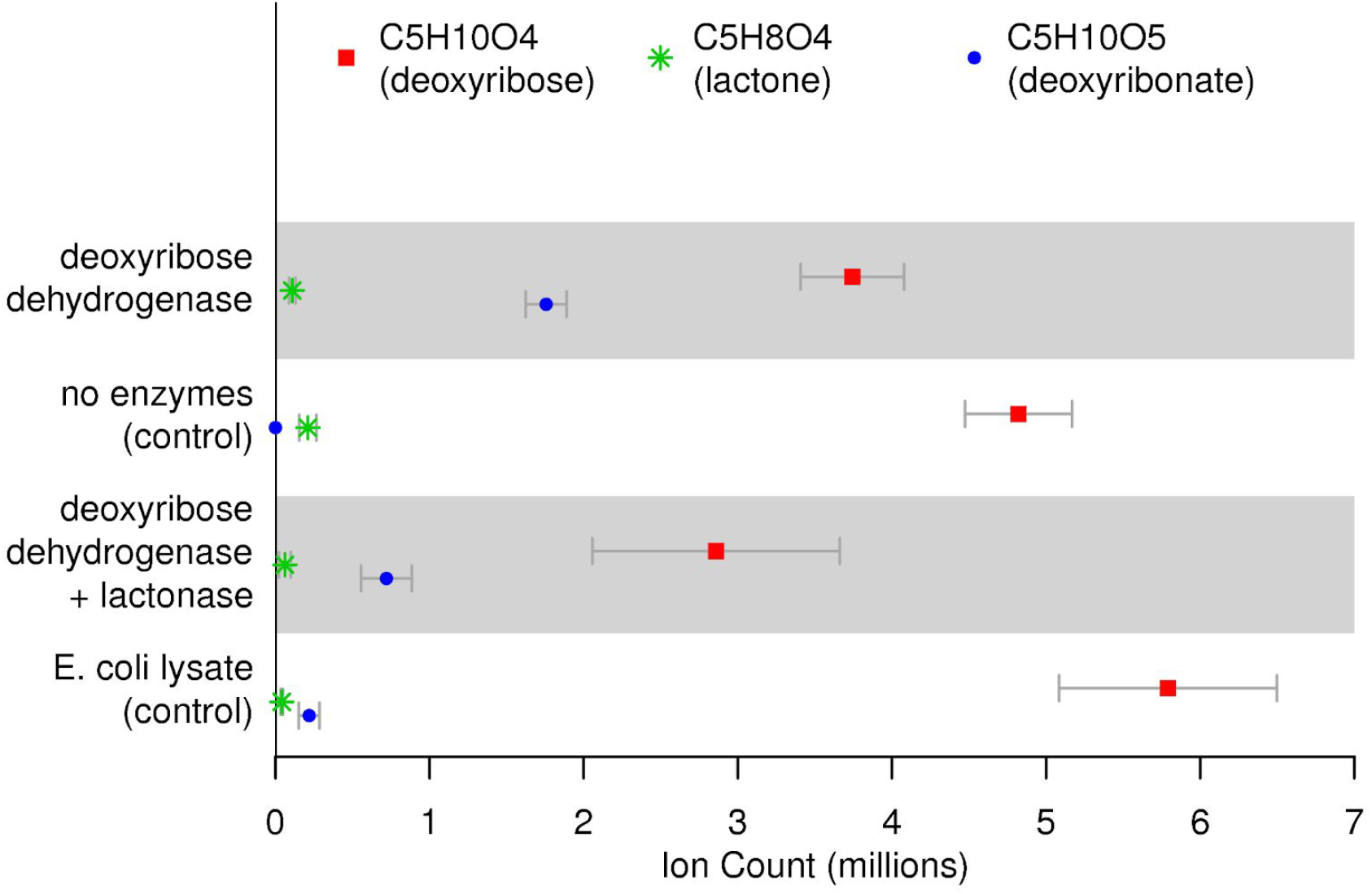
Reconstitution of the oxidation of deoxyribose to deoxyribonate *in vitro*. Deoxyribose, a potential deoxyribonolactone, and deoxyribonate were detected as sodium adducts (*m/z =* 157.05, 155.0, and 173.04, respectively). Each point is the average of 3 replicates and the error bar is the standard deviation.

**Figure S5:**
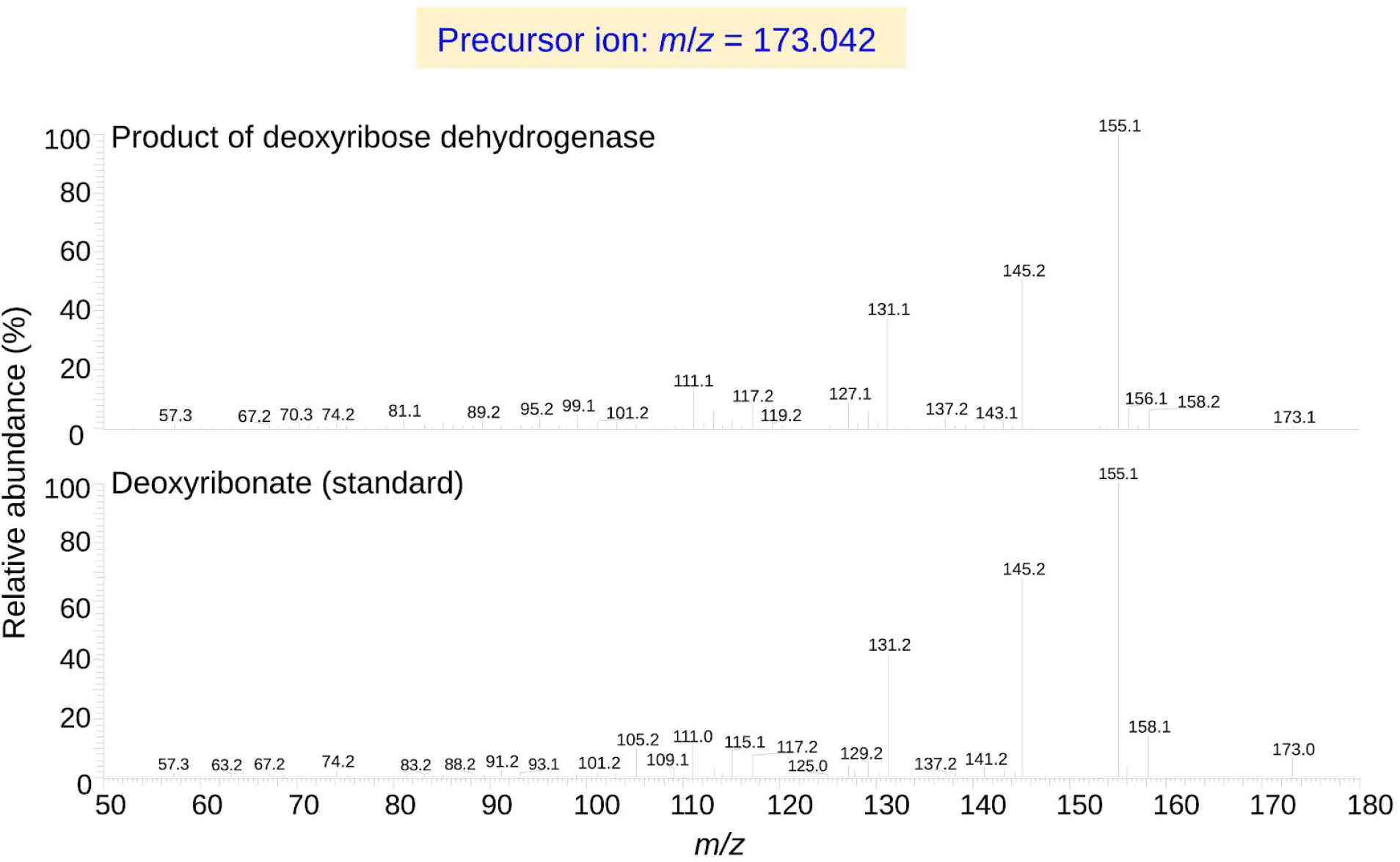
The tandem mass spectra of the product of deoxyribose dehydrogenase and of deoxyribonate.

**Figure S6:**
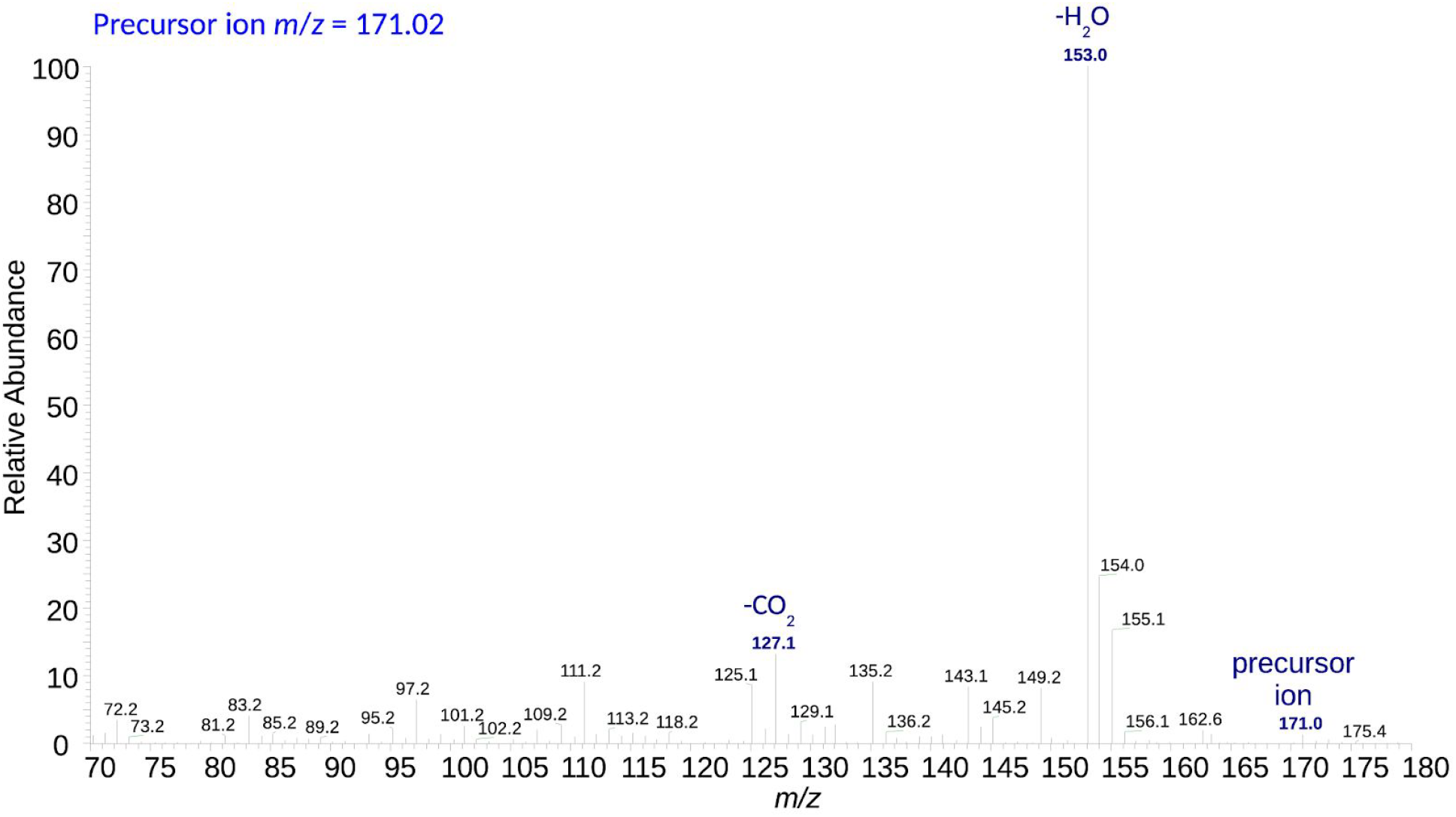
Tandem mass spectrum (MS/MS) of the putative 2-deoxy-3-keto-D-ribonate from deoxyribonate incubated with purified deoxyribonate dehydrogenase.

